# Multiome Perturb-seq unlocks scalable discovery of integrated perturbation effects on the transcriptome and epigenome

**DOI:** 10.1101/2024.07.26.605307

**Authors:** Eli Metzner, Kaden M. Southard, Thomas M. Norman

## Abstract

Single-cell CRISPR screens link genetic perturbations to transcriptional states, but high-throughput methods connecting these induced changes to their regulatory foundations are limited. Here we introduce Multiome Perturb-seq, extending single-cell CRISPR screens to simultaneously measure perturbation-induced changes in gene expression and chromatin accessibility. We apply Multiome Perturb-seq in a CRISPRi screen of 13 chromatin remodelers in human RPE-1 cells, achieving efficient assignment of sgRNA identities to single nuclei via an improved method for capturing barcode transcripts from nuclear RNA. We organize expression and accessibility measurements into coherent programs describing the integrated effects of perturbations on cell state, finding that *ARID1A* and *SUZ12* knockdowns induce programs enriched for developmental features. Pseudotime analysis of perturbations connects accessibility changes to changes in gene expression, highlighting the value of multimodal profiling. Overall, our method provides a scalable and simply implemented system to dissect the regulatory logic underpinning cell state.

## INTRODUCTION

Single-cell CRISPR screens have transformed the field of functional genomics by enabling simple, pooled experiments linking genetic perturbations to high-content phenotypes.^1, 2^ Perturb-seq, CROP-seq, and Mosaic-seq pioneered these approaches, expanding screen readouts from scalar measurements of cell fitness or reporter expression to comprehensive assessment of changes in transcriptional state.^3–6^ As single-cell assays have advanced, techniques profiling multiple modalities have integrated transcriptional data with relevant covariates,^7–10^ enabling new questions to be asked about the origins and regulation of cellular states. In particular, multiome assays that simultaneously measure gene expression and chromatin accessibility in the same cells have the potential to help explain how chromatin state influences and directs the transitions between transcriptional states.^11, 12^ During development, changes in accessibility have been shown to “foreshadow” gene expression changes and ultimately specify cell lineages.^11, 13–15^ Conversely, bulk co-profiling of expression and accessibility changes following chemical perturbations has suggested that certain pathways induce expression changes within open chromatin independent of remodeling, while others trigger global rewiring of chromatin state.^12, 16–18^ The unclear interplay between regulatory layers calls for a method to simultaneously characterize many perturbation-induced effects on the transcriptome and epigenome in parallel. These connections would have diverse applications because of their mechanistic interpretability, from elucidating regulators of cellular plasticity to gathering data to constrain causal modeling of gene regulatory networks.

To this end, we introduce Multiome Perturb-seq, an extension of the Perturb-seq platform that captures simultaneous scRNA-seq and scATAC-seq measurements of perturbation effects. Our protocol adapts the CROP-seq vector^5^ to be compatible with commercially available 10x Single Cell Multiome ATAC + Gene Expression reagents (“10x Multiome”). The core challenge is robust sgRNA assignment from nuclear cDNA, as isolating nuclei is a requirement of scATAC-seq protocols. To overcome this, we modified the CROP-seq vector and developed a new amplification protocol to enhance recovery of CROP-seq sgRNA identity transcripts (“IDs”), increasing perturbation assignment to 68%. We apply this protocol to a library of 13 chromatin remodelers from three regulatory complexes, demonstrate how Multiome Perturb-seq can profile and organize expression and accessibility changes into coherent effects on cell state, and interpret the dynamic regulatory processes connecting these two layers of gene regulation.

## RESULTS

### An optimized sgRNA assignment workflow recovers perturbation identities of single nuclei

We chose 10x Multiome as the target platform due to its ubiquity, quality, and the commercial availability of reagents, but this introduces a challenge for sgRNA capture: the corresponding beads contain oligo-dT primers to capture mRNA, and ATAC primers to capture transposed genomic DNA, but critically lack primers for capture of sgRNAs. This limitation restricted our ID expression and assignment strategy to the polyadenylated transcript-dependent CROP-seq approach, which, in the case of nuclei isolation, is hindered by active nuclear export of polyadenylated mRNAs.^19^ To mitigate this effect, we engineered a modified CROP-seq vector and library preparation protocol to increase capture and assignment of sgRNA ID transcripts **(Fig. 1a)**. We incorporated an internal high-activity T7 promoter^20^ upstream of the sgRNA cassette (similarly to methods used to boost sgRNA signal in spatial contexts^21^), enabling efficient linear amplification of IDs from cDNA before index PCR **(Methods, Document S2)**. We also flanked the ID transcript with short sequence elements (“SIRLOINs”) that increase nuclear localization of RNA,^22^ added a partial 10x TSO binding sequence to assist amplification of IDs in case of incomplete template switching during reverse transcription,^23^ and removed the highly structured woodchuck hepatitis virus post-transcriptional regulatory element (WPRE), which we reasoned might impair efficiency of reverse transcription. We observed that SIRLOIN elements drove a modest increase in nuclear abundance of IDs coupled with a modest decrease in overall expression, consistent with their original characterization **(Fig. S1a)**, and did not alter knockdown efficiency of cell surface markers **(Fig. S1b)**. Finally, we included a cassette for inserting auxiliary barcodes (e.g. as lineage markers^24, 25^), which can be read out alongside IDs via altered sequencing parameters in our protocol if desired.

**Fig. 1:**
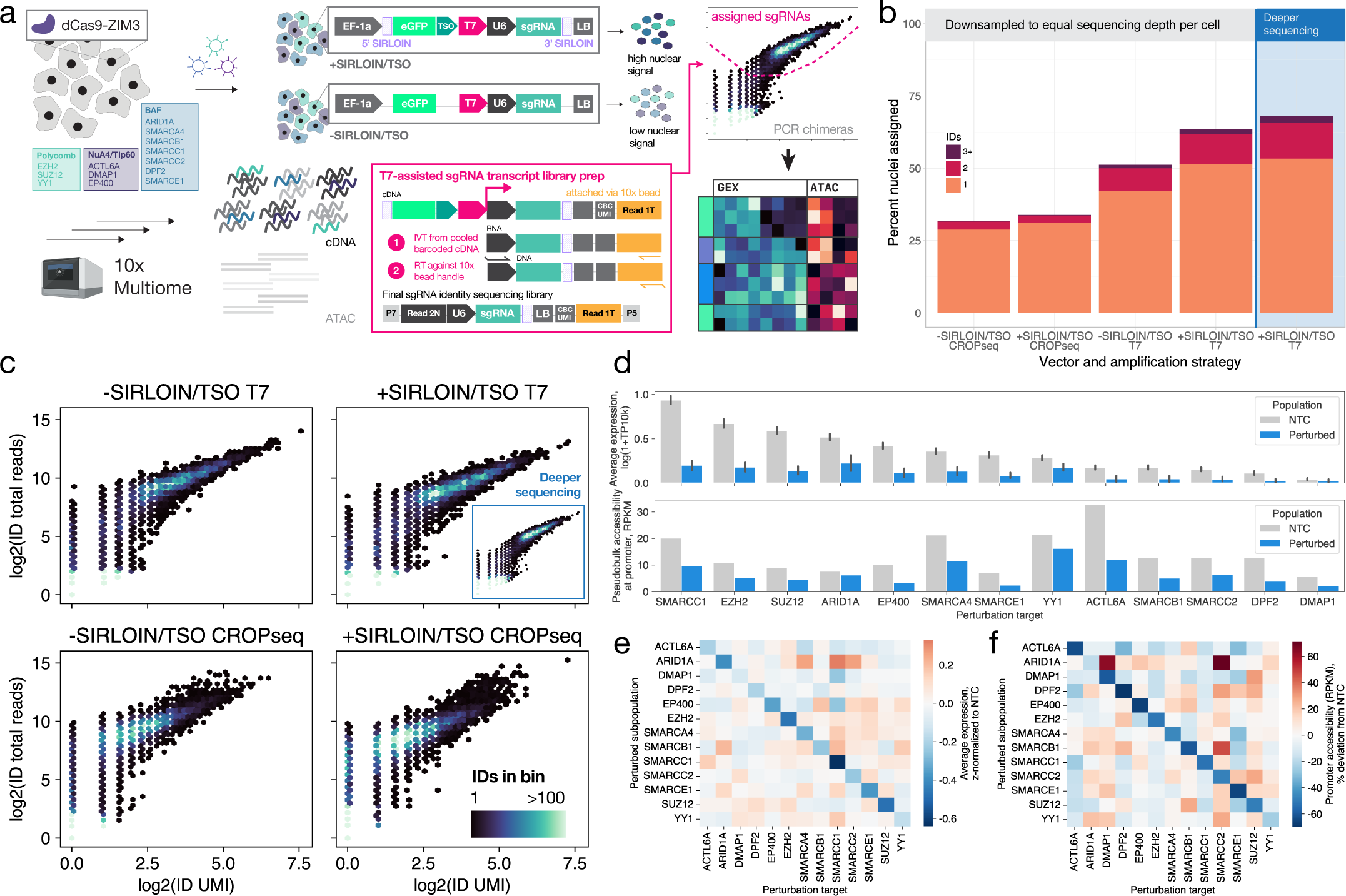
Improved perturbation assignment to single nuclei with T7-assisted sgRNA library prep and an engineered CROP-seq vector. **a)** Schematic of Multiome Perturb-seq experimental design, library, and guide assignment strategy. sgRNA identity transcripts are biased toward nuclear retention by addition of dual SIRLOIN elements flanking the sgRNA cassette (purple), and a partial 10x TSO capture sequence (green) assists recovery. Transcripts are linearly amplified via an internal high-activity T7 promoter from barcoded cDNA. U6: modified U6 Pol III promoter, LB: lineage barcode cassette. **b)** Percentage of single nuclei assigned to 1, 2, or 3+ sgRNA identities using traditional CROP-seq or T7-assisted library prep, in a standard CROP-seq vector with T7 promoter (-SIRLOIN/TSO) or a CROP-seq vector with T7 promoter, dual SIRLOINs and partial TSO capture sequence (+SIRLOIN/TSO), at indicated sequencing depths (downsampled for comparison, deeper sequencing for downstream analysis). **c)** Density of UMI and total sequencing reads of sgRNA identity transcripts amplified by traditional CROP-seq or T7-assisted library prep in nuclei-containing droplets, in each vector condition, downsampled to equal sequencing depth per nucleus. **d)** Average single-cell expression and pseudobulk promoter accessibility of perturbation target genes in NTC-assigned populations vs. populations assigned to each perturbation. Error bars for single-cell expression show 95% confidence intervals. **e)** Average normalized expression of perturbation target genes in perturbed subpopulations. **f)** Pseudobulk normalized accessibility at promoters of perturbation target genes in perturbed subpopulations.

We applied Multiome Perturb-seq in a CRISPRi screen of RPE-1 cells stably expressing ZIM3-dCas9^26^ using both the fully engineered Multiome Perturb-seq vector and a vector lacking SIRLOIN and TSO elements. Importantly, these hTERT-immortalized but untransformed cells allow us to investigate gene regulation in a more physiological context than cancer cell lines. Our library included three non-targeting control (NTC) sgRNAs and 13 sgRNAs targeting chromatin remodelers belonging to three regulatory complexes^27–30^ **(Fig. 1a, Supplementary Table 1)**. We isolated nuclei from RPE-1-ZIM3 populations expressing each version of the library, encapsulated them in two 10x Multiome lanes targeting 10,000 nuclei each, and recovered 9,318 (+SIRLOIN/TSO) and 9,504 (-SIRLOIN/TSO) nuclei after cellranger-arc cell calling **(Methods)**.

At equal sequencing depth of 850 reads per nucleus, traditional CROP-seq PCR-based library amplification yielded 31.8% assignment of single nuclei to at least one ID, improving to 33.8% in the +SIRLOIN/TSO condition **(Fig. 1b, Fig. S2)**. In contrast, our T7 amplification protocol yielded 51.2% assignment in the -SIRLOIN/TSO condition, increasing to 63.4% in the +SIRLOIN/TSO condition. Deeper sequencing of the T7 +SIRLOIN/TSO library to *∼*4,000 reads per nucleus modestly increased assignment to 68.0%. We traced these efficiency differences to shifts in library composition. Linear amplification of IDs before PCR shifted final ID UMI and total read density toward the high-UMI/high-read mode representing assignable transcripts, with a more pronounced shift in the +SIRLOIN/TSO condition **(Fig. 1c, Methods)**. Overall, our ID assignment workflow represents a greater than 40% improvement over the current state-of-the-art single-cell ATAC-seq screening approach on the 10x platform (Spear-ATAC: 48% assignment),^31^ while additionally providing gene expression phenotypes.

Analysis of the +SIRLOIN/TSO lane, after filtering for single ID assignments and performing basic quality control **(Fig. S3, Methods)**, yielded a final population of 4,724 nuclei with an average of 2,739 genes and 19,202 unique ATAC fragments detected. We observed robust on-target knockdown of expression and decreased accessibility at target gene promoters in perturbed populations **(Fig. 1d)**. Notably, we observed these effects across a range of native chromatin contexts, with both high-accessibility (e.g., *ACTL6A*) and low-accessibility (e.g., *DMAP1*) promoters displaying decreased accessibility upon perturbation. This result is consistent with the mechanism of the ZIM3 CRISPRi effector, whose KRAB domain recruits repressive chromatin-modifying complexes to induce heterochromatin formation.^26, 32^ While the on-target effect of each perturbation was strongest as expected, we additionally observed modest up-regulation of *SMARCA4*, *SMARCC1*, and *SMARCC2* in the sgARID1A-containing population, suggesting crosstalk between these BAF complex members **(Fig. 1e)**. Increased expression of *SMARCC2* was concordant with increased accessibility of the *SMARCC2* promoter in sgARID1A-containing cells **(Fig. 1f)**. Overall, our optimized assignment workflow efficiently associates perturbations with single-cell transcriptomes and accessibility profiles.

### Induced accessibility and expression changes provide complementary and largely non-overlapping descriptions of perturbation effects

We next analyzed the global phenotypes induced by perturbations. Identification of differentially expressed genes and differentially accessible peaks within each perturbation **(Supplementary Table 2, Methods)** revealed highly variable behaviors among individual perturbations: some mainly affected the chromatin landscape (*ARID1A*, *SMARCC2*), while others mostly modified gene expression (*DMAP1*, *EP400*), and some altered both equally (*SMARCE1*, *YY1*) **(Fig. 2a)**. In line with previous work,^16^ we found that while examples of concordant changes in accessibility and expression at loci other than perturbation targets were present in 7 of 13 perturbations, these instances were relatively rare, and most differential expression and accessibility events across perturbations were independent **(Supplementary Table 3)**.

**Fig. 2:**
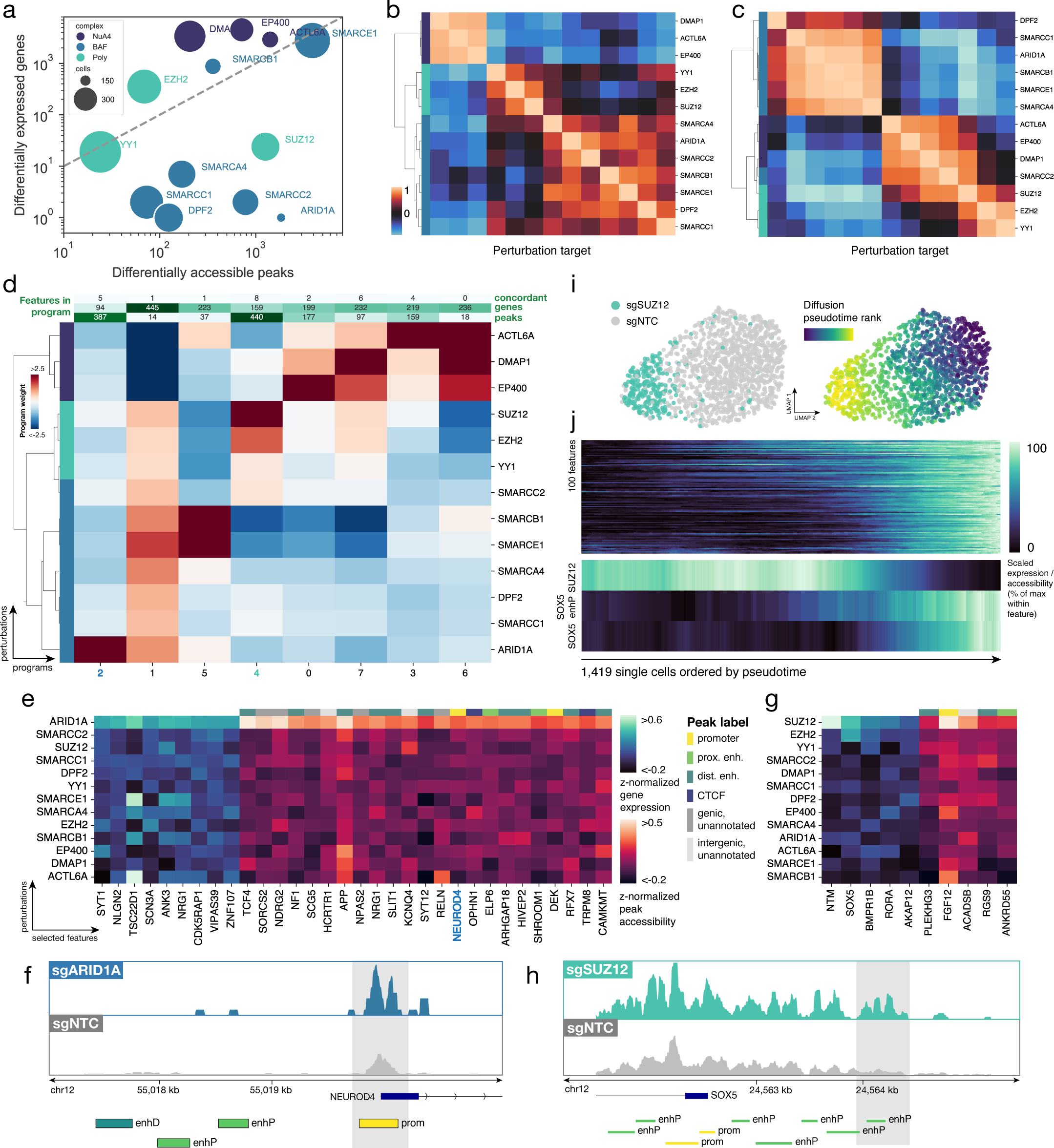
Multiome Perturb-seq characterizes transcriptional and epigenetic state to elucidate regulatory relationships. **a)** Number of differentially expressed genes (q < 0.1, Kolmogorov-Smirnov test on normalized expression, Benjamini-Hochberg FDR correction) vs. number of differentially accessible peaks (q < 0.1, Mann-Whitney test on paired insertion counts within peaks, BH FDR correction) for each perturbation target. Point sizes represent number of cells assigned each sgRNA identity. **b)** Hierarchical clustering of average PCA embeddings of normalized expression in perturbed subpopulations. **c)** Hierarchical clustering of average spectral embeddings of ATAC tile matrix in perturbed subpopulations. **d)** Program weights on perturbed subpopulations and number of features with nonzero loadings per program from sparse PCA embedding of 7,885 genes and 5,060 peaks differentially expressed or accessible in at least one perturbation condition. **e)** Normalized expression and accessibility of select genes related to nervous system development in Program 2; peaks labeled with nearest gene. **f)** Pseudobulk ATAC-seq track at the *NEUROD4* promoter in sgARID1A-containing and sgNTC-containing cells, with ENCODE cCRE annotations (prom, promoter; enhP, proximal enhancer; enhD, distal enhancer). Differential peak selected as feature in Program 2 highlighted in grey. **g)** Normalized expression and accessibility of *SOX5* and *SOX5* target genes in Program 4. **h)** Pseudobulk ATAC-seq track at the *SOX5* promoter for sgSUZ12-containing and sgNTC-containing populations, with ENCODE cCRE annotations (prom, promoter; enhP, proximal enhancer; enhD, distal enhancer). Differential peak in proximal enhancer highlighted in grey. **i)** UMAP representations of 1,419 sgSUZ12-containing and sgNTC-containing single cells, colored by sgRNA identity and diffusion pseudotime rank. **j)** Top) Expression and accessibility of top 100 Program 4 features by weight in the population shown in Fig. 2i. (Bottom) Expression of *SUZ12*, accessibility of the *SOX5* proximal enhancer shown in Fig. 2h, and expression of *SOX5* in the same population. Single cells are ordered along the x-axis by their diffusion pseudotime values, with each feature separately scaled to a range of 0-1.

To compare how the distinct readouts organized perturbations, we performed hierarchical clustering on both expression and accessibility profiles **(Methods)**. Both modalities separated targets into known complexes, with tighter clustering mirroring the relative impact of different complexes on either the transcriptome or chromatin state: NuA4 members clustered distinctly in transcriptome space, while BAF complex members were clearly delineated in accessibility space **(Fig. 2b, c)**. These differences underscore that expression and accessibility profiles offer complementary information, enabling nuanced characterization of perturbation effects across related targets.

### Co-embedding genes and peaks into sparse programs organizes effects on cell state

To organize the effects of perturbations into coherent programs, we concatenated filtered expression and accessibility matrices and co-embedded genes and peaks via non-negative sparse PCA **(Methods)**. This approach, applied to 7,885 genes and 5,060 peaks differentially expressed or accessible in at least one perturbation (and for peaks, limited to those within 50 kb of a gene), yielded 8 sparse programs, each containing hundreds of individual features **(Fig. 2d, Supplementary Table 4)**. Notably, programs varied in composition: some programs (2, 4) were largely peak-driven, others (1, 5, 6) gene-driven, and others (0, 3, 7) mixed. Labeling each peak with its nearest gene to facilitate interpretation, we focused on Programs 2 and 4, reasoning that these represented a “slow” and more consequential rewiring of cell state, in contrast to possibly noisier, “faster” gene-driven programs operating within open chromatin.

Program 2, which was highly specific to the sgARID1A-containing population, featured multiple regulators of neural development and function as top hits **(Fig. 2e)**, intriguing given the neuroectodermal ancestry of the retinal pigment epithelium.^33^ Further suggesting cellular plasticity induced by this program, we observed increased accessibility at the promoter of *NEUROD4*, a transcription factor normally expressed in retinal progenitors throughout embryonic development^34^ that specifies photoreceptor differentiation and function^35^ **(Fig. 2f)**. While *ARID1A* knockdown did not directly induce NEUROD4 expression, the increased promoter accessibility suggests it may be primed for activation by additional stimuli, hinting at a synergistic role for *ARID1A* and *NEUROD4* in cell fate determination.

### Ordering accessibility and expression changes along pseudotime trajectories elucidates regulatory relationships

Finally, the single-cell resolution of Multiome Perturb-seq has the potential to yield deeper insights into the relationship between chromatin accessibility and gene expression. In Program 4, specific to Polycomb members *EZH2* and *SUZ12*, we found high feature weights on *SOX5* (a TF implicated in nervous system and skeletal system development^36, 37^). The program contained several SOX5 target genes **(Fig. 2g)**, and we observed a global increase in accessibility around the *SOX5* TSS as well as a differentially accessible peak corresponding to a *SOX5* proximal enhancer in sgSUZ12-containing cells **(Fig. 2h)**.

Given these concordant changes, we reasoned that sgSUZ12-containing cells could anchor a pseudotime trajectory ordering heterogeneity within both control and perturbed populations, allowing us to visualize sequential chromatin opening preceding increased gene expression.^38–40^ As expected, a diffusion pseudotime metric constructed on a combined population of 1,419 sgSUZ12-containing and sgNTC-containing cells (Methods) mapped a trajectory from control cells to perturbed cells **(Fig. 2i)**. We visualized the expression and accessibility of the top 100 features by weight within Program 4 in single cells ordered by pseudotime, revealing broad upregulation aligned with this principal axis. **(Fig. 2j, top)** Some features “fired” earlier along the trajectory than others, suggesting candidate regulatory relationships between these “early” features and “late” targets. In the case of *SOX5*, accessibility of the proximal enhancer precedes increased *SOX5* expression and mirrors *SUZ12* knockdown, suggesting a causal role for *SUZ12* in repression of this locus and for this proximal enhancer in modulating the expression level of *SOX5* **(Fig. 2j, bottom)**. Overall, Multiome Perturb-seq’s unique combination of single-cell resolution, CRISPR perturbations, and multi-modal profiling underscores its utility as a tool for assembling regulatory networks and elucidating mechanistic relationships connecting transcriptional states to their epigenomic foundations.

## DISCUSSION

We developed Multiome Perturb-seq, a single-cell CRISPR screening method that efficiently assigns perturbation identities to simultaneous transcriptional and epigenomic phenotypes. Though we target use with robust, commercially available reagents, the protocol is likely simple to adapt to other workflows like combinatorial indexing^12, 41–43^ or to in vivo applications that require nuclei isolation. Our T7-assisted amplification protocol and engineered Multiome Perturb-seq vector recovered perturbation identities for 68% of input nuclei, a >40% improvement over comparable techniques for scATAC-seq.^31^ This method revealed that different perturbations distinctly alter chromatin accessibility and gene expression, underscoring the value of our multiomic approach in decoupling these layers of gene regulation. By leveraging single-cell heterogeneity, we could disentangle cis-regulatory dynamics and visualize sequential changes to chromatin state that foreshadowed changes in gene expression. Future screens employing our vector’s lineage barcodes and measuring changes over time could be used to study epigenetic heterogeneity and memory within clonal lineages.^44^

Looking forward, we envision that Multiome Perturb-seq will not only enable experiments that advance basic biological discovery but also facilitate rational engineering of cell state. By simultaneously profiling the transcriptome and epigenome, this method provides a comprehensive view of both a cell’s current state and potential future avenues for perturbation, as the epigenomic data can reveal genes newly primed for activation. Our observation of increased *NEUROD4* promoter accessibility in response to *ARID1A* knockdown exemplifies this concept, suggesting *NEUROD4* may be primed for activation and represent the next step in a defined regulatory path. Such insights could guide the design of synergistic perturbations, where an initial change in chromatin state sensitizes specific genes to subsequent interventions. This approach could direct programmed combinatorial genetic perturbations towards previously unreachable cell states, with relevance for both basic research and translational applications.

## Acknowledgements

We gratefully acknowledge all the members of the Norman Lab for their valuable thoughts and discussion on this work. We thank Duaa H. Al-Rawi for assistance with nuclei isolation. This work was funded by NIH Director’s New Innovator Award DP2 GM140925 (to T.M.N.). This work was funded in part through the NIH/NCI Cancer Center Support Grant P30 CA008748. We acknowledge the use of the MSKCC Integrated Genomics Operation and Flow Cytometry Cores, funded by the NCI Cancer Center Support Grant (CCSG, P30 CA008748), Cycle for Survival, and the Marie-Josée and Henry R. Kravis Center for Molecular Oncology.

## Author Contributions

Conceptualization, E.M., K.M.S. and T.M.N. Methodology, E.M., K.M.S., and T.M.N. Formal analysis, E.M. Investigation, E.M. and K.M.S. Writing – Original Draft, E.M. Writing – Review and Editing, E.M., K.M.S., and T.M.N. Visualization, E.M. Supervision, T.M.N. Funding Acquisition, T.M.N.

## Supplemental Information

**Document S1:** Figures S1-S3 and Supplemental Figure legends.

**Document S2:** Protocol for T7-assisted CROP-seq library prep, related to Figure 1.

**Table S1:** sgRNA protospacers used in this study, related to Figure 1 and Figure 2.

**Table S2:** Differentially expressed genes and ATAC-seq peaks in each perturbation condition, related to Figure 2.

**Table S3:** Quantification of concordant changes in differential expression and accessibility, related to Figure 2.

**Table S4:** Annotated features and weights for sparse programs, related to Figure 2.

**Table S5:** Oligonucleotide sequences used in this study.

## METHODS

### Experimental

#### Construction of engineered CROP-seq vectors

Vector pEM040 (+SIRLOIN/TSO) was constructed sequentially from vector pKS010 (available upon request from lead contact). A high-activity T7 promoter with appropriate overlaps was ordered from IDT as a duplexed oligo (Table S5) and cloned into PpuMI-linearized pKS010 via HiFi assembly (NEB #E2621). Ultramer oligos containing the SIRLOIN consensus sequence^22^ and appropriate HiFi assembly overlaps were ordered from IDT (Table S5) and sequentially cloned into the 5’ and 3’ UTRs of the CROP-seq transcript, first via linearization with BsiWI (5’ UTR) to create an intermediate vector, then via double digestion with SalI and BamHI (3’ UTR), removing an internal Nextera Read 2 from the lineage barcode cassette in the parent vector. The partial TSO sequence was ordered from IDT as an Ultramer oligo and similarly cloned by HiFi assembly into an intermediate vector linearized with MluI. Vector pEM047 (-SIRLOIN/TSO) was constructed by adding the T7 promoter as described above and digesting with SalI / BamHI to remove the Nextera Read 2, but the 3’ SIRLOIN oligo was replaced with a duplex oligo ligated into the digested product with Hi-T4 ligase (NEB #M2622) at RT for 1 hr. For each cloning reaction, assembled vectors were transformed into NEB Stable E. coli, (NEB #C3040H) and grown at 32°C overnight. Single colonies were picked and miniprepped (Qiagen #27104), and sequences were verified by Plasmidsaurus whole plasmid sequencing.

#### sgRNA library construction and quality control

Vectors pEM040 and pEM047 were sequentially digested with BstXI and BlpI, and the desired 8119 bp / 7973 bp products were gel purified from an 0.8% agarose gel. Each vector contains a 1133-bp LacZ fragment that is removed by double digestion with BstXI and BlpI, allowing visualization of complete digestion during gel purification. sgRNAs were chosen based on previous published and unpublished data demonstrating on-target activity, and NTC guides were selected that minimized differential expression in previous screens. sgRNA libraries were ordered as IDT oPools with 5’ overhang 5’-agtatcccttggagaaccaccttgttg-3’ and 3’ overhang 5’-gtttaagagctaagctggaaacagcatagcaag-3’. For library cloning, oPools were resuspended at 100 ng/µL, and 10 µL (1 µg) was used as input to a second strand synthesis reaction with 2x KAPA HiFi Master Mix (Roche, #07958935001) and an equimolar amount of primer P276R (Table S5). The reaction was run in a thermal cycler with the following parameters: 95° 3’, 64° 15” (ramp down 0.1°C/sec), 72° 10’, 4° hold. After purification of the double-stranded product (NEB #T1030), the library was cloned into digested pEM040 and pEM047 via HiFi assembly (NEB #E2621) for 1 hr at 50°C, using a 5:1 molar ratio of insert library to vector. Electroporation was performed with an Eppendorf Eporator into Endura Electrocompetent Cells (LGC 60242-2) per manufacturer protocol. After electroporation, each reaction was incubated in 1 mL Recovery Media at 32°C for 1.5 hr, then expanded into 100 mL LB + carbenicillin for overnight culture. Plasmid libraries were produced via midiprep (Qiagen #12941).

For QC of the constructed libraries, sgRNA protospacers were amplified via 2x KAPA HiFi Master Mix (Roche, #07958935001) from midipreps with primers containing Illumina P5 and P7 adapters and Nextera i7 indices (oJR232, oJR233, Table S5). Following 2x ProNex purification (Promega NG2001), amplicons were sequenced on an Illumina MiSeq with PE70 read structure. Resulting fastq files were processed with a custom Python script to quantify representation of each sgRNA in the library (Fig. S3).

#### Cell lines and lentiviral packaging

RPE-1-ZIM3 cells were obtained from Replogle and Saunders et al., 2022^45^ and maintained in DMEM:F12 media (Gibco, 11320033), 10% Fetal Bovine Serum (FBS) (VWR, 97068-085), 100U penicillin and 100 mg/mL streptomycin (Gibco, 15140122), and 0.01 mg/mL Hygromycin B (Thermo Fisher, 10687-010). LentiX 293T cells were purchased from Takara Bio (632180) and maintained in DMEM (Gibco #11965092), 10% Fetal Bovine Serum (FBS) (VWR, 97068-085), 100U penicillin and 100 mg/mL streptomycin (Gibco, 15140122). For lentiviral packaging, LentiX cells were grown to 70-80% confluence and co-transfected with library vector, transfer, and packaging plasmids psPAX2 and pMD2G using TransIT-LTI Transfection Reagent (Mirus, MIR 2300). 24 hr post-transfection, media was changed to DMEM + 10% BSA (Sigma #A9418). Virus was harvested by filtering collected media through a 0.45µm filter 48 hr post-transfection.

#### Multiome Perturb-seq: experimental procedure

For the Multiome Perturb-seq experiment, populations of 1.2 *×* 10^7^ RPE-1-ZIM3 cells were infected with lentivirus containing each library at a concentration of 400 µL harvested lentivirus per million cells (targeting MOI 0.1) and plated into 3 T175 flasks each in the presence of 8 µg/mL polybrene (Day 0). On Day 3, cells were sorted for GFP fluorescence by FACS (FacsSymphony S6, BD Biosciences). Infection titers of 6.6% and 9.4% were obtained for the +SIRLOIN/TSO vector and -SIRLOIN/TSO vector, respectively. 1.2 *×* 10^6^ and 2.0 *×* 10^6^ cells were recovered from sorting, and each population was plated in a 10cm dish. On Day 6, 1 *×* 10^6^ cells from each population were prepared for encapsulation via nuclei isolation, following the 10x protocol “Nuclei Isolation for Single Cell Multiome ATAC + Gene Expression Sequencing” (CG000365, Rev C) with 4 min lysis time (Step 2d) and verification of complete lysis by Trypan Blue staining and visualization on the Countess II FL (Life Technologies). We observed that digitonin quality is extremely important to high-quality nuclei extraction in RPE-1 cells; digitonin that is expired or not thawed with rapid mixing at 65°C before use will result in incomplete lysis and affect the quality of extracted nuclei.

Isolated nuclei were resuspended in 500 µL PBS + 1 U/µL Sigma Protector RNase inhibitor (Sigma #3335402001), and 25 µL 7-AAD (Thermo #00-6993-50) was added to stain intact nuclei. Samples were immediately sorted by FACS (FacsSymphony S6, BD) to purify 7-AAD+ nuclei and remove debris, resulting in yields of 409,591 nuclei (+SIRLOIN/TSO) and 404,204 nuclei (-SIRLOIN/TSO). While sorting nuclei is an optional step of the 10x Multiome protocol, we observed that sorting for intact nuclei reduced ambient RNA which could impede ID assignment. Sorted nuclei were resuspended in 10x Nuclei Buffer targeting a concentration of 4,000 nuclei/µL, assuming 30% loss of the sorted yield during resuspension. Targeting 10,000 nuclei per reaction and accounting for the 10x-indicated recovery factor of 1.61, we loaded 4 µL ((10, 000 nuclei *×* 1.61)/(4, 000 nuclei/µL)) of each nuclei stock into transposition reactions per 10x protocols (Chromium Next GEM Single Cell Multiome ATAC + Gene Expression User Guide, Rev F), and proceeded with GEM generation, barcoding, and cleanup using the Chromium Controller and Chromium Next Gen Chip J (10x Genomics #1000283) with no protocol modifications. ATAC libraries were constructed with no protocol modifications and 7 PCR cycles for ATAC library amplification (Step 5.1d). cDNA amplification was performed with no protocol modifications and 6 PCR cycles (Step 6.1d). Gene expression libraries were constructed with no protocol modifications, using 12 PCR cycles for library amplification (Step 7.5d). After PicoGreen quantification and quality control by Agilent TapeStation, libraries were pooled to equimolar concentrations and sequenced to a target depth of 20,000 reads per cell on an Illumina NovaSeq X in a PE28/88 run using the NovaSeq X 10B or 25B Reagent Kit (100 or 300 Cycles) (Gene Expression) or to a target depth of 25,000 reads per cell on a NovaSeq 6000 in a PE50 run using the NovaSeq 6000 S1, S2, or S4 Reagent Kit (100 or 200 Cycles) (ATAC). The loading concentration was 1 nM (NovaSeq X) or 0.6-1.2 nM (NovaSeq 6000) and a 1% spike-in of PhiX was added to each run for quality control purposes.

#### Multiome Perturb-seq: T7-assisted library prep

*A full step-by-step protocol is available in Document S2.* To construct perturbation identity transcript libraries from pooled cDNA, 10 µL (25% of final yield by volume) of amplified cDNA from each library (Chromium Next GEM Single Cell Multiome ATAC + Gene Expression User Guide Rev F, Step 6.2n product) was in-vitro transcribed (IVT) with the NEB Quick T7 HiScribe Kit (NEB #E2050) in 25 µL reactions per manufacturer protocol. IVT reactions were incubated at 37°C for 2 hr. 3.8 µL dH2O and 1.2 µL DNase I were then added directly to each reaction (30 µL total reaction volume), and the reactions were incubated at 37°C for 15 min. IVT RNA product was purified with 1.8x RNAClean (Beckman Coulter #A63987) and eluted in 16.5 µL RNase-free dH2O. 1 µL product was used to quantify RNA yield with the Qubit RNA BR kit. The entire purified IVT product was used as template for reverse transcription with the Maxima H Minus First Strand cDNA Synthesis Kit (Thermo # K1652) as follows. On ice, 2 µL 10 µM RT primer (T7-TruSeqRead1, Table S5) and 1.25 µL 10 mM dNTPs were added to 15.5 µL IVT product. The reactions were incubated at 65°C for 5 min and immediately returned to ice. To each reaction on ice, 5 µL 5x RT buffer and 1.25 µL RT Enzyme Mix were added. Reactions were incubated at 50°C for 30 min and heat-inactivated at 85°C for 5 min. Next, the entire RT product was used directly in index PCR across 5 50 µL reactions to ensure that the volume of unpurified RT product did not exceed 10% of each individual PCR reaction. A 5x PCR master mix for each sample was built on ice as follows: to 25 µL unpurified RT product, we added 97 µL dH2O, 1.5 µL 100 µM P065-P5 (P5 reverse primer), 1.5 µL 100 uM P065-N7xx (indexed P7 forward primer, captures U6 promoter directly 5’ of protospacer), and 125 µL 2x KAPA HiFi Master Mix (Roche #07958935001) (Table S5). The master mix was divided among 5 50-µL reactions in a PCR strip, and PCR was performed with the following parameters: 95° 3’, 8 cycles of [98° 15”, 70° 20”], 72° 1’, 4° hold. PCR reactions were pooled and purified via 1.15x ProNex (+SIRLOIN/TSO) or 1.2x ProNex (-SIRLOIN/TSO). Yield was measured with the Qubit dsDNA HS kit (Thermo #Q32851) and amplicons were visualized as single peaks at 584 and 516 bp respectively on a Bioanalyzer (Agilent) using the High Sensitivity DNA Chip. After PicoGreen quantification and quality control by Agilent TapeStation, libraries were pooled to equimolar concentrations and sequenced to a target depth of 3,000 reads per cell on an Illumina NovaSeq X in a PE28/88 run using the NovaSeq X 10B or 25B Reagent Kit (100 or 300 Cycles). The loading concentration was 1 nM and a 1% spike-in of PhiX was added to the run for quality control purposes.

#### Multiome Perturb-seq: PCR enrichment CROP-seq library prep

ID transcripts were isolated from Gene Expression library (Chromium Next GEM Single Cell Multiome ATAC + Gene Expression User Guide Rev F, Step 7.6q product) for comparison with T7-assisted library prep by traditional CROP-seq. Transcripts were amplified from 200 ng library product across 5 50-µL PCR reactions with 2x KAPA HiFi Master Mix (Roche, #07958935001), P065-P5, and P065-N7xx (Table S5). PCR was performed with the following parameters: 95° 3’, 20 cycles of [98° 15”, 70° 10”], 72° 1’, 4° hold. Reactions were pooled and purified with 1.1x (+SIRLOIN/TSO) or 1.15x (-SIRLOIN/TSO) ProNex. We reduced ProNex ratios slightly vs. T7 library prep to reduce the presence of starting material in final libraries, a known issue with CROP-seq. Yield was measured with the Qubit dsDNA HS kit (Thermo #Q32851) and amplicons were visualized as single peaks at 584 and 516 bp respectively on a Bioanalyzer (Agilent) using the High Sensitivity DNA Chip. After PicoGreen quantification and quality control by Agilent TapeStation, libraries were pooled to equimolar concentrations and sequenced on a NovaSeq X in a PE28/88 run using the NovaSeq X 10B or 25B Reagent Kit (100 or 300 Cycles). The loading concentration was 1nM and a 1% spike-in of PhiX was added to the run for quality control purposes. While we targeted the same sequencing depth as the T7 libraries, the presence of gene expression library starting material in the final CROP-seq library resulted in fewer ID reads at the same total sequencing depth. Of 30 million target reads, we recovered 8.33 million (+SIRLOIN/TSO) and 13.8 million (-SIRLOIN/TSO).

#### RT-qPCR characterization of CROP-seq vectors

Leftover sorted nuclei and whole cells from the Multiome Perturb-seq experiment were used to quantify ID transcript expression. Total RNA was extracted from each sample (+/- SIRLOIN/TSO, nuclei and whole cells, 4 samples total) with the NEB Total RNA Miniprep kit (NEB #T2010). 10 ng nuclear RNA and 100 ng whole-cell RNA was used as template for RT-qPCR using the NEB Luna Universal One-Step RT-qPCR kit (NEB #E3005). PCR was performed per manufacturer recommendations, though the RT temperature was raised to 60°C. We targeted two 250 bp amplicons in the ID transcript for consistency, one within the coding sequence of GFP and the other within the U6 promoter. The *MALAT1* nuclear RNA was used as a normalization control, similarly to the original characterization of SIRLOIN elements.^22^ C_t_ values were measured on a CFX96 Real-Time System (Bio-Rad), and ΔC_t_ was calculated as the difference between amplicon and *MALAT1* C_t_ values in each condition. ΔΔC_t_ was calculated by subtracting the C_t_ values of the +SIRLOIN/TSO samples from that of the -SIRLOIN/TSO samples, and expression fold change for each amplicon in the +SIRLOIN/TSO condition was calculated as 2*^−^*^ΔΔCt^. Standard errors were propagated through C_t_ calculations as previously described.^46^

#### Flow cytometry validation of on-target knockdown

1.5 *×* 10^5^ RPE-1-ZIM3 cells were transduced with lentivirus containing CD81 or NTC sgRNAs in +SIRLOIN/TSO and -SIRLOIN/TSO CROP-seq vectors to a target MOI of 0.1 via plating with 8 µg/ml polybrene (Day 0). Transduced cells were sorted for GFP positivity on Day 3. On Day 6, cells were harvested, washed with PBS, and resuspended in 180 µL staining buffer (BD #554657). 20 µL CD81-APCH7 antibody (BD #656154) was added, and cells were incubated protected from light at 4°C for 15 min. Cells were then washed with 1 ml PBS and resuspended in 200 µL staining buffer, and CD81 signal was analyzed on an Attune CytPix flow cytometer (Thermo).

### Computational

#### sgRNA identity calling

For analysis, sgRNA assignments were called on the T7 +SIRLOIN/TSO library with deeper sequencing using a custom Python script, cropseq_call_guides_multiome.py. Briefly, cellranger-7.1.0 was run on ID library reads, associating sgRNA-containing reads with 10x cell barcodes and UMIs (cellranger count --chemistry=ARC-v1). Based on cells called by cellranger-arc (below), reads were filtered to those associated with cell-containing droplets and aligned to the protospacer library with fuzzy string matching (rapidfuzz). Reads were collapsed by cell barcode and UMI, creating a dataframe of the ID transcripts in each cell (one of 16 options based on alignment to the protospacer library) labeled with UMI and total read counts. The relationship between UMI and total read counts was used to separate true ID transcripts from PCR artifacts as follows. First, UMI and total read counts were log2-transformed, and possible “Top IDs” representing true assignments with log2(UMI) >= 2.75 were selected. An HDBSCAN model (min_samples = 10, metric = ‘l1’, cluster_selection_epsilon = 0.5) was fit to Top IDs to further filter these putative true assignments. Minority clusters and IDs categorized as noise (label -1) were removed, and the cluster with higher average UMI was selected as representative of true assignments. A logistic regression model was trained on all Top IDs, then applied to the entire dataset of ID transcripts to infer membership in the true assignment cluster. IDs for which the model predicted membership in this cluster were kept as true assignments, and other IDs were removed. This procedure results in a clear plane of separation between called and non-called transcripts (Fig. S2), and removes likely PCR chimeras (e.g., transcripts for which the UMI-total read relationship deviates significantly from confident assignments, such as transcripts with a single UMI and read representing a likely template-switching event during index PCR). We additionally filtered the model predictions such that no ID transcript with less than 3 UMI would be called, regardless of its cluster prediction.

For comparisons of assignment between library prep and vector conditions, we downsampled individual reads from the above dataframe of ID transcripts (pd.DataFrame.sample()) before collapsing to UMI and total read counts, such that the total number of reads in this dataframe divided by the number of cells called by cellranger was constant (850 reads per cell, based on the minimum common sequencing depth available across conditions). We ran the ID assignment script as above, but changed the “Top ID” log2(UMI) threshold to 2 (T7) or 1.75 (CROPseq).

#### Single-cell data processing and QC filtering

cellranger-arc-2.0.2 (10x Genomics) was used for alignment of Gene Expression and ATAC libraries to the transcriptome and genome respectively (reference: cellranger-arc-GRCh38-2020-A-2.0.0), using cellranger-arc count with default parameters. For gene expression processing, sgRNA ID assignments were merged with cellranger gene expression outputs in Scanpy,^47^ and quality control metrics were calculated with sc.pp.calculate_qc_metrics(). We inspected these metrics by sgRNA assignment status (Fig. S3) and filtered low-quality cells based on total UMI and library complexity (number of unique genes). Mitochondrial genes were removed, and cells were filtered to those with single sgRNA assignments. The resulting AnnData object was further filtered using the perturbseq codebase^3^ to remove genes expressed in fewer than 5% of cells with strip_low_expression(threshold = 0.05), and counts were then z-normalized to control by normalizing total counts in all cells to median UMI, subtracting the NTC mean of each gene, and dividing by the NTC standard deviation. The resulting normalized matrix was read back into Scanpy to regress out cell cycle genes (obtained from Harmonizome.^48^)

For ATAC processing, 10x fragment files were converted to tile matrices with SnapATAC2.^49^ QC metrics were calculated, and cells with ≤1000 unique fragments were removed. From the genome-wide tile matrix, 2.5 *×* 10^5^ features were selected for downstream analyses based on accessibility in all cells. The combined quality control process applied to both modalities left 4,724 high-quality nuclei with single sgRNA assignments and complete expression and accessibility profiles.

#### Identification of differentially expressed genes and differentially accessible peaks

Differentially expressed genes were called on the filtered z-normalized matrix described above with a Kolmogorov-Smirnov test (perturbseq.ks_de()). P-values were corrected for false discovery rate via the Benjamini-Hochberg procedure, and genes with FDR-corrected q < 0.1 were considered differentially expressed. Differentially accessible ATAC peaks were identified using SnapATAC2’s suggested workflow. From the filtered tile matrix described above, peaks were called with MACS3^50^ (snap.tl.macs3()) on pseudobulk accessibility profiles quantified with paired-insertion counting^51, 52^ for each perturbation target and NTC, and a cell-by-peak matrix with fixed-width, non-overlapping features was created with snap.tl.merge_peaks() followed by snap.tl.make_peak_matrix(counting_strategy = ‘paired-insertion’). For each perturbation, differentially accessible peaks were identified with a Mann-Whitney test (scipy.stats) between counts in NTC and perturbed cells, performed on each feature of the cell-by-peak matrix for which any perturbed or NTC cells had at least one count. P-values were corrected for false discovery rate via the Benjamini-Hochberg procedure, and peaks with FDR-corrected q < 0.1 were considered differentially accessible. Log fold change values were calculated as the log2-transformed ratio of mean counts in perturbed cells to control cells. A pseudocount of 1 *×* 10*^−^*^3^ was added to the numerator and denominator during this calculation to avoid infinite values. To identify concordant differential changes, each peak was labeled with its nearest gene, and matches were considered concordant if both the peak and its nearest gene were differentially expressed or accessible with q < 0.1 and the sign of the gene expression z-score was equal to the sign of the peak log fold change.

#### Annotation of ATAC-seq peaks with ENCODE cis-regulatory elements

A .bed file of cis-regulatory elements was downloaded from ENCODE SCREEN.^53^ To label ATAC differential features, PyRanges was used to intersect this bed file with the coordinates of fixed-width merged peaks composing the features of the cell-by-peak matrix described above. Peaks were labeled with CREs based on overlap of at least 1bp. To label total peaks for Table S3, unique peaks from MACS3 output were intersected and labeled with the regulatory element for which the peak had the largest overlap. For a finer-grained characterization of promoters, we supplemented promoter annotations with a database of human promoters downloaded from the Eukaryotic Promoter Database,^54^ and we extended coordinates by 950 bp upstream to represent promoter regions. Features annotated as “prom” overlap a promoter region. The SOX5 enhancer peak shown in Figs. 2i and 2j was labeled as overlapping a promoter region, but contained a proximal enhancer in the ENCODE dataset. We characterize the peak as an enhancer based on its ENCODE annotation to maintain consistency, but the feature remains labeled as overlapping a promoter region in the raw data (e.g., in Table S4).

#### Clustering of expression and accessibility profiles

Gene expression profiles were clustered by decomposing the processed single-cell expression matrix with PCA into 10 components. Component values were grouped by perturbation identity and averaged, and these pseudobulk embeddings were hierarchically clustered using 1 - Pearson correlation as the distance metric and Ward’s linkage. ATAC profiles were clustered by decomposing the processed single-cell tile matrix with SnapATAC2’s spectral embedding algorithm into 11 components. We removed the first component of this embedding, finding that it correlated strongly with the number of unique fragments recovered in each single cell. The remaining 10 components were similarly grouped by perturbation identity and averaged, and the pseudobulk spectral embeddings were clustered by hierarchical clustering with 1 - cosine similarity as the distance metric and Ward’s linkage. For clustermap generation in each case, we used hc.linkage(method = ‘ward’, optimal_ordering = True) from scipy.cluster.hierarchy.

#### Sparse PCA co-embedding of expression and accessibility features

To organize perturbation effects and decompose peak-driven vs. gene-driven changes, we adapted a sparse PCA approach used by our group to interpret Perturb-seq data to the Multiome setting. For this analysis, we first z-normalized paired insertion counts within the peak matrix as follows: from the cell-peak matrix described above, features were pre-filtered to remove the lower 25% based on count values in all cells and any peaks more than 50 kb away from the nearest gene. Means and standard deviations of counts in NTC cells for each feature were then generated, and the filtered peak matrix was z-normalized by subtracting the NTC mean and dividing by the NTC standard deviation. From the z-normalized peak matrix, we then removed peaks that were not differentially accessible in any perturbations, resulting in 5,060 peaks. We similarly z-normalized gene expression matrices as described above, and removed genes not differentially expressed in any perturbations. This resulted in a concatenated expression/accessibility matrix of 5,060 peaks and 7,885 genes for downstream analysis.

Our sparse PCA model was constructed by modifying sklearn’s SparsePCA object to constrain feature weights to be nonnegative, using dict_learning(positive_code = True). This procedure promotes interpretability by 1) limiting the number of gene and peak features within each resulting program, as the model is fit with a regularization term penalizing nonzero feature weights; and 2) ensuring that features within a program are co-regulated in the same direction by each perturbation, via the nonnegativity constraint. We fit a sparse PCA model to the above concatenated matrix, deciding empirically on the number of programs to fit (limited by the low-rank structure of the matrix, with only 13 perturbations and 3 underlying complexes) and the value of the regularization parameter alpha. Fitting 8 programs with alpha = 0.25 resulted in expected clustering of perturbation weight vectors into known complexes, with a sparsity level that allowed for substantial gene-peak heterogeneity between programs while not sacrificing the value of the method as a feature selection tool. Peak features in each program were then labeled with their nearest gene and ENCODE cis-regulatory element annotations based on overlap, if applicable. Features, weights, and annotations for each program are available in Table S4. Nervous system genes highlighted in Fig. 2 were manually curated from Program 2 features. *SOX5* target genes were obtained from Harmonizome.^48^

#### Generation and visualization of ATAC-seq coverage tracks

Bigwig coverage tracks were generated at pseudobulk level for each perturbation and NTC cells as follows. The .bam output of cellranger-arc was filtered with pysam to include only reads with the ‘CB’ tag corresponding to perturbation-assigned cell barcodes, creating a pseudobulk .bam file for each perturbed subpopulation and NTC. These files were indexed with samtools index and converted to bigwig coverage tracks with deepTools bamCoverage --normalizeUsing RPGC --effectiveGenomeSize 2913022398 --binSize 20. Tracks were visualized with pyGenomeTracks.^55^

#### Construction of pseudotime trajectories

We applied a diffusion pseudotime approach as previously described^40^ to create an ordered embedding of control and perturbed single cells. We constructed a new gene expression matrix in scanpy by normalizing each cell to equal total UMI and log1p-transforming normalized counts, avoiding z-normalization to maintain meaningful values for NTC cells. Similarly, starting with the cell-by-peak matrix as above, we filtered peaks more than 50 kb away from the nearest gene and tf-idf normalized raw paired insertion count values. We then constructed an individual pseudotime trajectory for each perturbation as follows. We selected only features that were differentially expressed or accessible in the perturbation and concatenated these features into a combined expression and accessibility matrix. For many perturbations, the target gene and promoter were differentially expressed and accessible (Fig. 1); we removed these genes and peaks from the calculation to maintain on-target expression and accessibility as a “positive control” to evaluate the fidelity of our embedding in separating perturbed and control cells. We then scaled each feature to zero mean and unit variance and decomposed the matrix into 10 principal components. We constructed the nearest-neighbors graph on this PCA embedding using sc.pp.neighbors(metric = ‘correlation’), and constructed a diffusion map with sc.tl.diffmap(). To set a root cell for diffusion pseudotime construction (required by scanpy’s implementation), we identified the diffusion component with the highest absolute-value Pearson correlation with a binary perturbed/control variable for each single cell, and chose the cell at the appropriate extreme of this component (lowest value if correlation was negative, highest value if positive) as the root cell.

We then constructed the pseudotime with sc.tl.dpt(), and visualized cells ordered by their pseudotime rank via a UMAP embedding fit on the above concatenated differential expression / accessibility matrix. Pseudotime ranks aligned well with perturbed/control identities for most perturbations. Notably, alternative trajectories that do not require the a priori choice of a root cell (single-component spectral embedding, linear discriminant analysis) produced indistinguishable results when applied to the combined population. Visualizations (Fig. 2j) were constructed by smoothing ordered single-cell expression and accessibility values with a 500-cell rolling average, then scaling each feature separately to a range of 0-1.

---

**Fig. S1:**
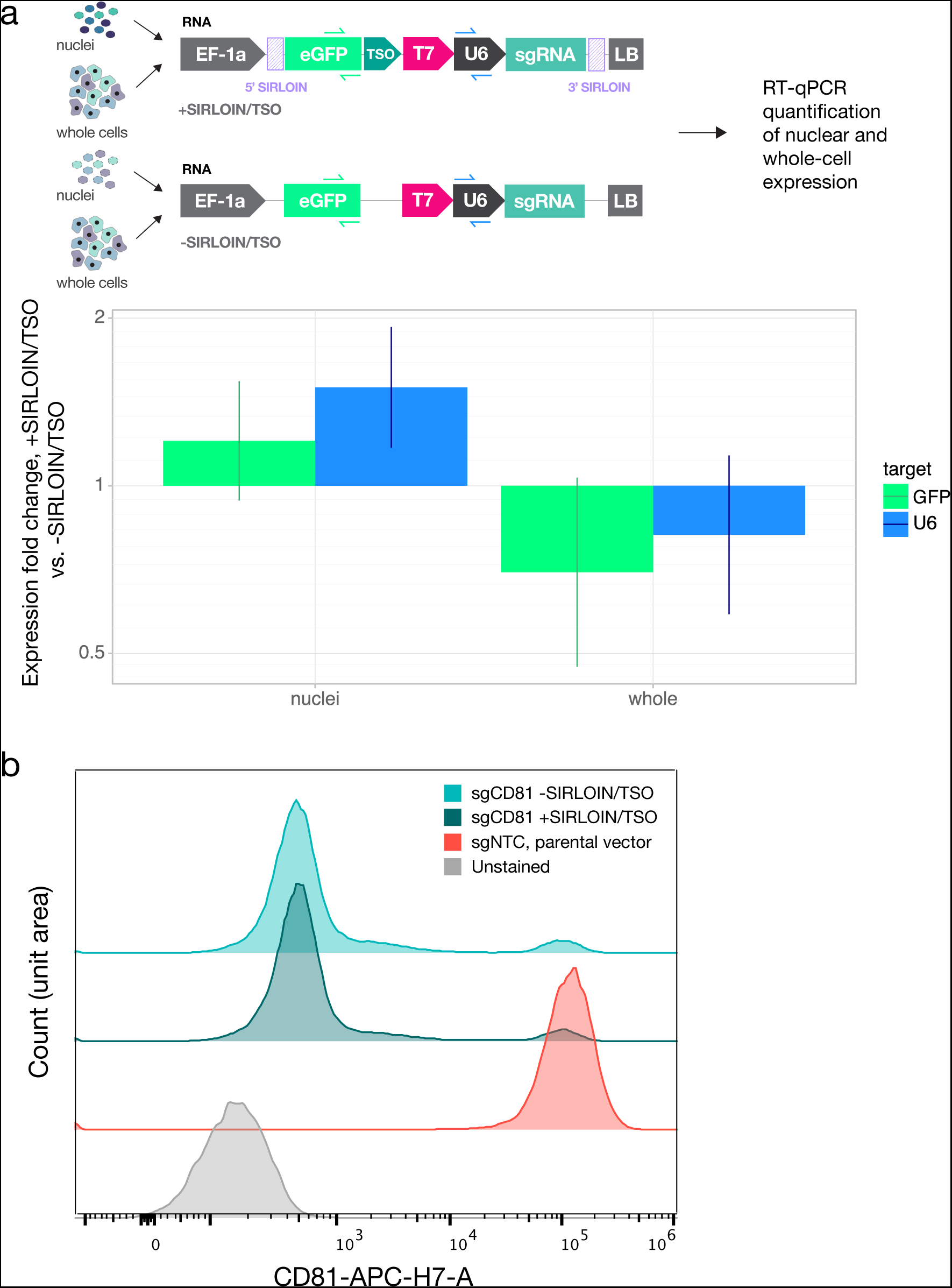
Characterization of engineered CROP-seq vectors. **a)** RT-qPCR quantification of nuclear and whole-cell expression of CROP-seq vectors with and without SIRLOIN elements (purple). Error bars show 95% confidence intervals across 3 technical replicates of each condition targeting a unique amplicon within the ID transcript, each of length 250 bp. **b)** Flow cytometry validation of on-target knockdown of cell surface marker CD81 by sgRNAs expressed from each vector in RPE-1-ZIM3 cells.

**Fig. S2:**
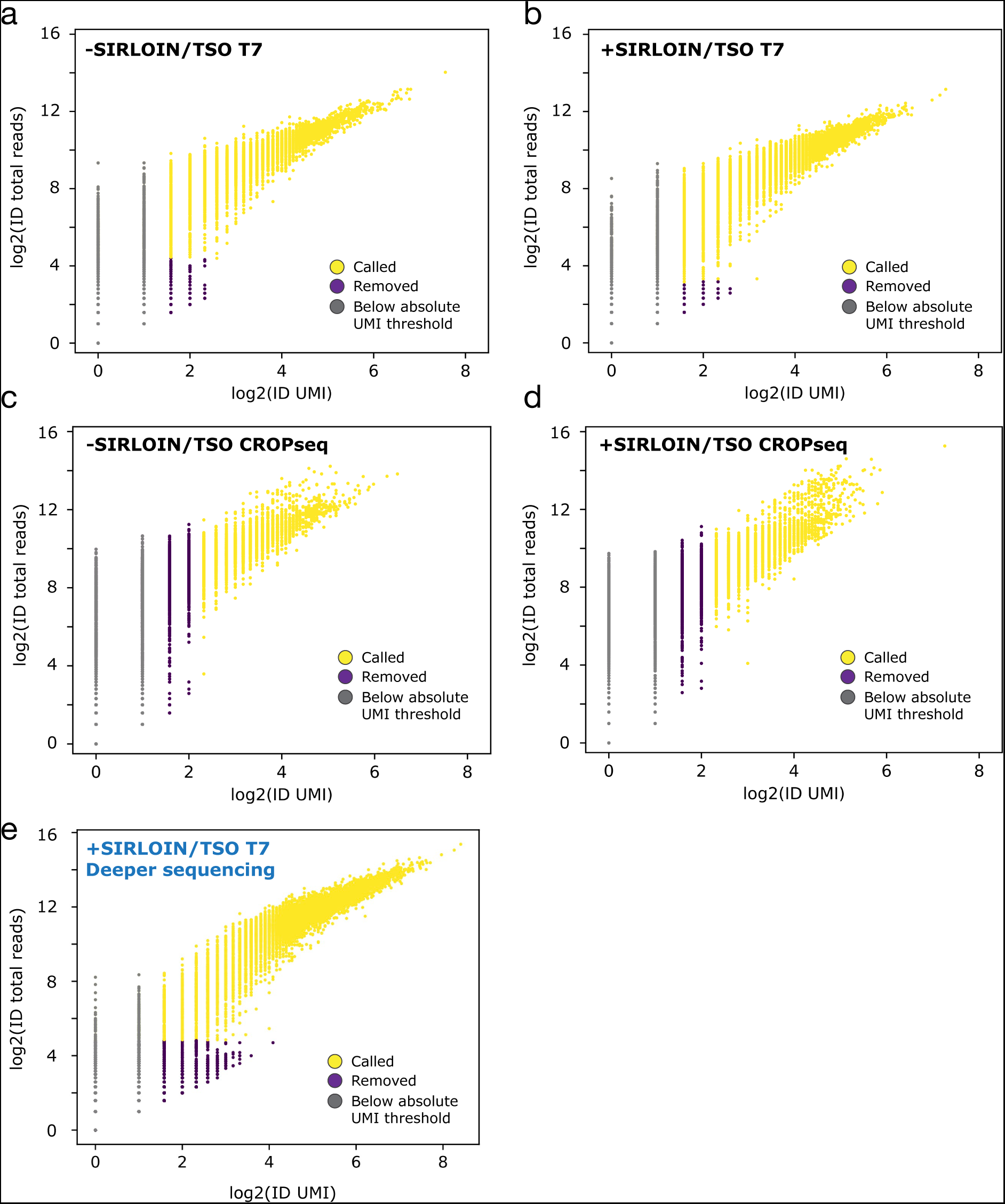
Outputs of CROP-seq ID calling pipeline. **a)** Called vs. non-called ID transcripts in nuclei-containing droplets, -SIRLOIN/TSO vector, T7-assisted library prep, downsampled to 850 reads per nucleus. **b)** Called vs. non-called ID transcripts in nuclei-containing droplets, +SIRLOIN/TSO vector, T7-assisted library prep, downsampled to 850 reads per nucleus. **c)** Called vs. non-called ID transcripts in nuclei-containing droplets, -SIRLOIN/TSO vector, CROPseq library prep, downsampled to 850 reads per nucleus. **d)** Called vs. non-called ID transcripts in nuclei-containing droplets, +SIRLOIN/TSO vector, CROPseq library prep, downsampled to 850 reads per nucleus. **e)** Called vs. non-called ID transcripts in nuclei-containing droplets, +SIRLOIN/TSO vector, T7-assisted library prep, full sequencing depth (4,034 reads per nucleus).

**Fig. S3:**
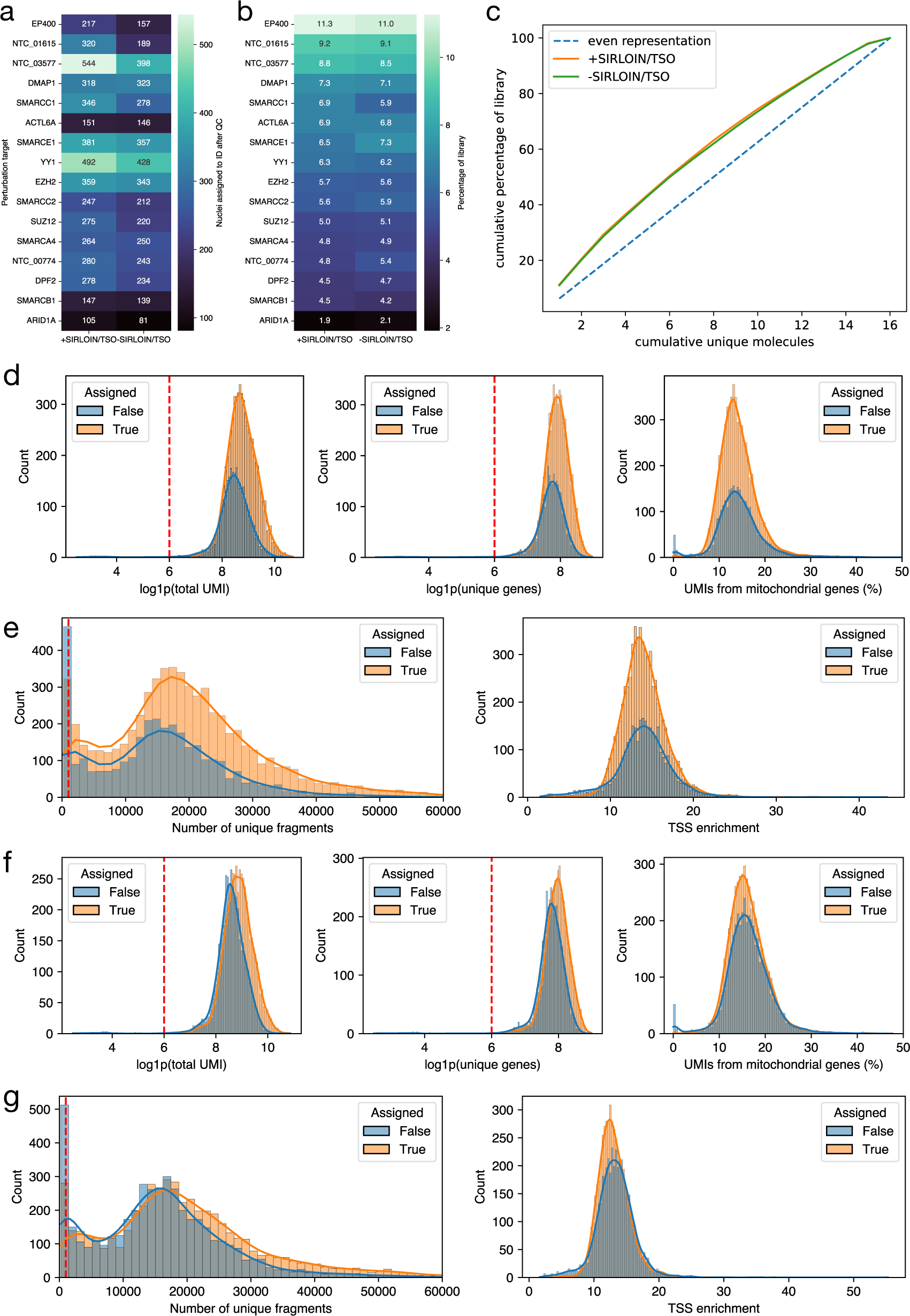
Quality control metrics related to the Multiome Perturb-seq screen. **a)** Number of cells assigned to each perturbation in each vector condition after QC. **b)** Representation of sgRNAs in cloned libraries. **c)** Gini plot showing skew of sgRNA representation in each library. **d)** Gene expression UMI, library complexity (number of unique genes), and mitochondrial transcript percentage in all cellranger-called cells, stratified by ID assignment, in +SIRLOIN/TSO condition. Dashed lines show QC filters applied. **e)** Unique ATAC fragments recovered and TSS enrichment in all cellranger-called cells, stratified by ID assignment, in +SIRLOIN/TSO condition. Dashed lines show QC filters. **f)** Gene expression UMI, library complexity (number of genes), and mitochondrial transcript percentage in all cellranger-called cells, stratified by ID assignment, in -SIRLOIN/TSO condition. **g)** Unique ATAC fragments recovered and TSS

