## Supplementary material for "Multiome Perturb-seq unlocks scalable discovery of integrated perturbation effects on the transcriptome and epigenome": Document S2

### Document S2 - Protocol for T7-assisted sgRNA ID library prep

This protocol involves manipulation of nanogram amounts of RNA: before starting, clean benchtop (preferably with an RNase-decontaminating solution such as RNase-away) and use new RNase-free dH2O and pipette tips.

#### 1) T7 amplification of ID transcripts from amplified cDNA and final ATAC library

1. Start with amplified cDNA product from 10x Multiome Protocol Step 6.2n; carry through 25% of product by volume (10 out of 40 µL) to T7 amplification, regardless of yield, but note that the maximum input to each T7 reaction is 1 µg template DNA
2. Assemble the T7 reaction at RT as follows, using the NEB Quick T7 HiScribe Kit:

##### T7 amplification

**One reaction per sample:**

- 10 µL Step 6.2n cDNA
- 12.5 µL NTP buffer mix
- 2.5 µL T7 mix

3. Incubate at 37˚C for 2 hr
4. Add 3.8 ul dH2O, then 1.2 µL DNase I (included in T7 kit) to each reaction; total volume is now 30 µL
5. Incubate at 37˚C for 15 min (take out RNAClean beads to equilibrate to room temp now)
6. Clean up reactions with 1.8x RNAClean (54 µL beads / 30 µL sample); elute RNA in 16.5 µL dH2O
    a. Add 54 µL beads to 30 µL reaction; pipette mix 15x
    b. Incubate at RT for 5 min, then place on the 10x magnet (high) for 2 min
    c. Remove supernatant
    d. Wash 3x with 200 µL 70% EtOH (wait 30 sec between washes)
    e. Centrifuge briefly, remove remaining EtOH, air dry beads 10 min (they will look very dry but this is not an issue as in SPRI)
    f. Add 16.5 µL RNase-free dH2O directly to the beads; pipette mix 15x
    g. Incubate at RT for 2 min
    h. Place on the 10x magnet (low) for 2 min, then transfer 16.5 µL IVT product to a new tube strip
7. Measure yield on Qubit with RNA BR kit (if yield is too low for detection on RNA BR kit, proceed anyway; there is still RNA in the sample)

#### 2) Reverse transcription of T7-amplified RNA

1. Start with purified IVT product from Step 1.6 above; for each RT reaction, the maximum input of RNA is 500 ng per 20 µL reaction, but reactions can be scaled based on Step 1 yield
2. Pre-set a thermal cycler to 65˚C
3. Assemble the RT reaction on ice as follows, using the Thermo Maxima H Minus First Strand cDNA Synthesis Kit:

##### Reverse transcription

**One reaction per sample, volumes can be scaled based on Step 1 RNA yield:**

- 15.5 µL IVT product (Step 1.6 pdt)
- 2 µL 10 µM RT primer (T7_TruSeqRead1, Table S5)
- 1.25 µL 10 mM dNTPs

4. Incubate at 65˚C for 5 min, then immediately place on ice

5. To each reaction on ice, add 5 µL 5x RT buffer, then add 1.25 µL RT Enzyme Mix (volumes per 25 µL final reaction volume, can be scaled up)
6. Incubate at 50˚C for 30 min, then inactivate at 85˚C for 5 min

#### 3) Index PCR

1. Use entire unpurified RT product directly in index PCR, but volume of unpurified RT product should not exceed 10% of each PCR reaction
2. Primer sequences are available in Table S5; this index PCR adds a unique i7 index only to the library amplicon
3. For each sample, divide 25 µL unpurified RT product to 5 50 µL PCR reactions by making the following master mix and dividing among 5 tubes of a PCR strip; then run the PCR reaction with the following program:

##### Index PCR

**5x master mix per sample:**

- 25 µL RT pdt
- 1.5 µL 100 uM P065-N7xx
- 1.5 µL 100 uM P065-P5
- 125 µL 2x KAPA HiFi HotStart ReadyMix
- 97 µL dH2O

**Program:**

95˚ 3'
---8x---
98˚ 15"
70˚ 20"
---8x---
72˚ 1'
4˚ hold

4. Pool 5 PCR reactions per sample into 1.5ml Eppendorf tubes, and purify via ProNex, eluting in 50 µL; for the +SIRLOIN/TSO vector, the expected amplicon size is 584 bp, so we use a 1.15x ratio
    a. Equilibrate ProNex beads to RT and vortex thoroughly; add 288 µL beads (1.15x) to 250 µL sample
    b. Incubate at RT for 10 min, then place on an Eppendorf-compatible magnet for 2 min
    c. Remove supernatant
    d. Wash 2x with 1 mL wash buffer (wait 30 sec between adding and removing wash buffer)
    e. Remove remaining wash buffer, then air dry 5 min (beads can optionally be dried for longer than 5 min)
    f. Resuspend beads in 50.5 µL elution buffer
    g. Incubate 5 min, then place back on the magnet for 1 min
    h. Transfer 50 µL sample to a new tube strip

#### 4) Library construction QC

1. Measure yield of above Step 3.4 product on Qubit with dsDNA HS kit; calculate library molarity with Qubit mass and appropriate amplicon size
    a. Yield expected around 50-100 ng for nuclei, 100-200 ng for whole cells
2. Run appropriate dilution (e.g. 1:10 for 2-3 ng/µL Qubit undiluted product) on Bioanalyzer; a successful library prep is visualized as a single peak at 584 bp
3. Keep libraries at -20˚C for long-term storage

#### 5) Sequencing parameters

1. sgRNAs can be sequenced with standard 10x Gene Expression sequencing parameters (paired end 28/10/10/90): Read 1 captures the cell barcode and UMI, and Read 2 captures the sgRNA protospacer

2. For ID assignment, target sequencing depth is 3,000 read pairs per nucleus (e.g., 30 million paired-end reads for a sample in which 10,000 nuclei were targeted as input)

#### Example Bioanalyzer trace of final library

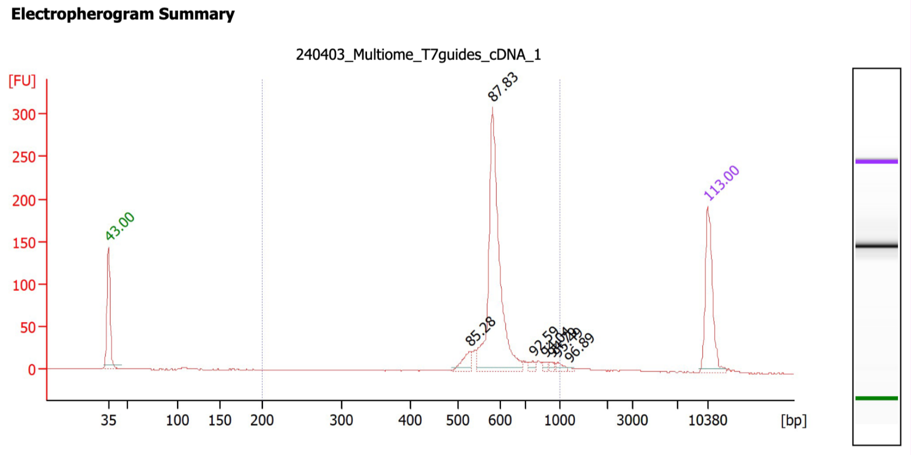


#### 6) Optional inclusion of lineage barcodes

This is an additional capability of the vector that is only lightly tested; the below methods may require more optimization.

1. Lineage barcodes are cloned into the vector via a protocol similar to sgRNA cloning. (Methods) We use a 16-bp barcode with 5' overhang 5'-AGTTCCCAAGATTCAACCGCGGAGGTCGAC-3' and 3' overhang 5'-GGATCCACAATCGCCAGTGCATAGCTGAC-3' (these are the "cloning spacers" in the below figure). The combined barcode and overhang oligos (5'overhang-barcode-3'overhang, 75 bp total) are similarly ordered as oPools and second strand synthesis is performed, and the double-stranded product is inserted via HiFi assembly into a SalI/BamHI double digest of the vector.
2. To sequence the barcode and sgRNA on the same run (desirable as IDs would only have to be assigned once), the barcode can be read during the i5 index read (since the T7 library prep protocol indexes samples with i7 only)
3. This would require a custom index primer as shown below; due to sequencer-level differences in i5 index read orientation, this strategy is only compatible with NextSeq/NovaSeq v1.5+ workflows.

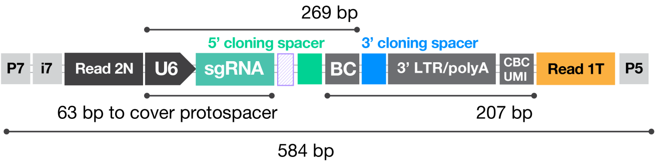


**Standard 10x Gene Expression parameters:**

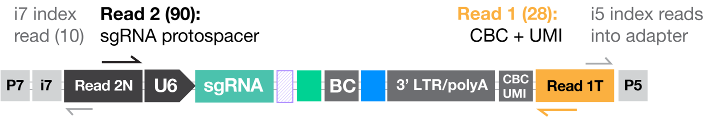


**Alternative parameters to sequence lineage barcodes:**

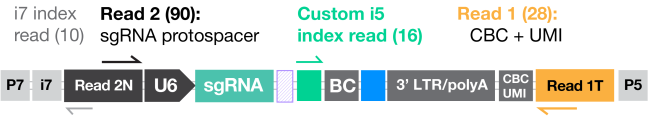
